# Astrocyte-expressed STAT3 regulates glutamate homeostasis and binge ethanol drinking in mice

**DOI:** 10.64898/2026.08.11.744063

**Authors:** Milagros Galan-Llario, Hu Chen, Emily Legge, Chloe M. Erikson, Roman Vlkolinsky, Jonathas Almeida, Michal Bajo, Marisa Roberto, Amy W. Lasek

## Abstract

Astrocytes play an important role in neuronal health. A critical function of astrocytes is to clear excess extracellular glutamate and prevent excitotoxicity. STAT3 is a transcription factor that promotes astrocyte development and astrocyte reactivity in neurodegenerative diseases and following central nervous system injury. To determine the innate molecular and behavioral functions of adult astrocyte-expressed STAT3 in a non-pathological state, we created conditional *Stat3* astrocyte knockout mice (Stat3 aKO) using Stat3^flox^ and the tamoxifen-activated Cre line, Aldh1l1-Cre/ERT2. We measured transcript levels of *Gfap*, a known STAT3 target gene, and glutamate transporter genes in the medial prefrontal cortex (PFC) of Stat3 aKO. *Gfap*, *Slc1a2* and *Slc17a8* transcripts were decreased in the PFC of Stat3 aKO of both sexes. GLT-1 protein, encoded by *Slc1a2*, was also reduced in the PFC of male Stat3 aKO. We recorded spontaneous excitatory post-synaptic currents (sEPSCs) in male Stat3 aKO and control prelimbic pyramidal neurons and found increased sEPSC amplitude, consistent with a hyper-glutamatergic state due to impaired glutamate clearance. To determine the behavioral consequences of STAT3 depletion in astrocytes, Stat3 aKO were tested for locomotor activity, anxiety-like behavior and binge ethanol consumption, behaviors linked to dysregulation of glutamate homeostasis. Stat3 aKO mice did not differ in locomotor activity or anxiety-like behavior; however, male Stat3 aKO mice consumed significantly less ethanol than controls. These results indicate that STAT3 in adult astrocytes is crucial for maintaining glutamate transporter levels in the adult brain and that astrocytic STAT3 promotes ethanol consumption in male mice.

**Main points:**

- *Gfap*, *Slc1a2* and *Slc17a8* expression are lower in the cortex of *Stat3* astrocyte knockout mice (Stat3 aKO)
- GLT-1 protein is decreased and glutamate neurotransmission is elevated in the cortex of male Stat3 aKO
- Male Stat3 aKO consume less ethanol

## 1. Introduction

Glutamate is the primary excitatory neurotransmitter in the mammalian central nervous system. Its extracellular concentration must be tightly regulated to enable proper synaptic transmission and prevent excitotoxicity. Astrocytes regulate and respond to extracellular glutamate levels via the Na⁺-dependent glutamate transporters GLT-1 (also known as EAAT2, encoded by *Slc1a2*) and GLAST (also known as EAAT1 encoded by *Slc1a3*). In addition, astrocytes also express the cystine/glutamate antiporter system xCT (encoded by *Slc7a11*), which mediates the non-vesicular release of glutamate into the extracellular space in exchange for cystine (Lewerenz et al., 2013). Among these, GLT-1 is the predominant transporter in the forebrain, responsible for the vast majority of extracellular glutamate uptake (Danbolt, 2001). Mice that globally lack GLT-1 develop lethal spontaneous seizures, while transgenic mice that overexpress GLT-1 show a higher seizure threshold than wild-type mice, underscoring the critical role of this transporter in preventing neuronal hyperexcitability (Peterson & Binder, 2020; Tanaka et al., 1997). Approximately 80% of GLT-1 is expressed by astrocytes (Furness et al., 2008). Like constitutive *Slc1a2* knockout, mice with conditional *Slc1a2* knockout in astrocytes exhibit increased mortality and more electrical seizure events as measured by electroencephalography (EEG), whereas knockout of *Slc1a2* in neurons does not affect lifespan or EEG measures (Petr et al., 2015), demonstrating the overall importance of astrocyte-expressed GLT-1 in regulation of glutamate homeostasis and brain health.

Signal Transducer and Activator of Transcription 3 (STAT3) is a transcription factor that regulates astrocyte development (Bonni et al., 1997; He et al., 2005). It is activated by phosphorylation of tyrosine 705 by Janus kinases (JAKs) in response to a wide array of cytokines, growth factors, and inflammatory mediators. Levels of tyrosine 705 phosphorylated STAT3 (pySTAT3) are increased in astrocytes in numerous disease states, such as spinal cord injury (Anderson et al., 2016; Herrmann et al., 2008; Okada et al., 2006), neurodegenerative diseases (Abjean et al., 2023; Ben Haim et al., 2015; Ceyzeriat et al., 2018; Hagemann et al., 2023; Kushwaha, Molesworth, Makarava, & Baskakov, 2025; Reichenbach et al., 2019) and alcohol addiction (Chen et al., 2021). STAT3 drives the expression of key astrocyte reactivity markers such as the intermediate filament protein GFAP, whose upregulation is a signature of a reactive astrocyte phenotype (Ceyzeriat, Abjean, Carrillo-de Sauvage, Ben Haim, & Escartin, 2016; Escartin et al., 2021). STAT3 also regulates expression of glutamate transporters in astrocytes in response to neurotoxic agents. For example, pySTAT3 was increased in mouse hippocampal astrocytes during methamphetamine withdrawal, along with decreased GLT-1 and GLAST protein, and methamphetamine withdrawal was associated with impaired glutamate clearance and disrupted spatial memory (Shi et al., 2021). Astrocyte-selective STAT3 knockdown in the hippocampus restored normal GLT-1 expression, glutamate clearance, and rescued the methamphetamine withdrawal-induced memory deficit (Shi et al., 2021), suggesting that STAT3 represses GLT-1 expression during methamphetamine withdrawal. In contrast, exposure of postnatal day 7 rats to the inhaled anesthetic sevoflurane resulted in decreased pySTAT3, GFAP, and GLAST in the hippocampus, an effect that was mimicked by treatment with a JAK inhibitor (Wang, Lu, Feng, & Zhang, 2016). These findings establish astrocytic STAT3 as either a repressor or activator of glutamate transporter expression and clearance capacity under neurotoxic conditions. Less is known regarding the function of astrocyte-expressed STAT3 under non-pathological conditions in adult animals.

To determine the innate function of STAT3 in astrocytes in adult mice, we deleted STAT3 from astrocytes using the Aldh1l1-Cre/ERT2 line crossed with Stat3^flox^ mice and treated 7-week-old mice with tamoxifen to induce Cre recombinase activity. We measured prefrontal cortex (PFC) expression of the well-established astrocytic STAT3 target gene *Gfap*, along with the glutamate transporter genes *Slc1a2* (GLT-1), *Slc1a3* (GLAST), *Slc7a11* (xCT) and *Slc17a8* (VGLUT3), which is involved in Ca^++^-dependent glutamate release from astrocytes (Ni & Parpura, 2009; Ormel, Stensrud, Chaudhry, & Gundersen, 2012). We also determined the effect of astrocyte *Stat3* deletion on glutamatergic neurotransmission in the PFC. Finally, we characterized the behavioral effects of *Stat3* deletion in astrocytes.

In addition to testing the *Stat3* astrocyte knockout mice for basal locomotor and anxiety-like behaviors, we also tested mice for several alcohol-associated behaviors, including ethanol consumption, because we previously demonstrated that mice treated systemically with a STAT3 inhibitor consumed less ethanol than vehicle-treated control mice in a binge ethanol consumption test (Hamada et al., 2021) and astrocytes have been shown to play a role in ethanol consumption (Erickson et al., 2021; Guizzetti et al., 2026; Kastner-Blasczyk, Hester, Reasons, Scofield, & Woodward, 2025; Tan & Ding, 2026). Decreased GLT-1 expression has also been associated with increased ethanol drinking (Lee et al., 2013; Tan, McBride, & Ding, 2026). Our results demonstrate that STAT3 in astrocytes is important for maintaining cortical GLT-1 levels in adult mice under non-pathological conditions and promoting binge alcohol consumption in male mice.

## 2. Methods

### 2.1 Animals

*Stat3* astrocyte knockout (Stat3 aKO) mice were generated by crossing Aldh1l1-Cre/ERT2 (B6N.FVB-Tg[Aldh1l1-cre/ERT2]1Khakh/J, JAX strain #031008) (Srinivasan et al., 2016) with homozygous Stat3^flox^ mice (B6.129S1-Stat3^tm1Xyfu^/J, JAX strain #016923)(Moh et al., 2007). Aldh1l1-Cre/ERT2 and Stat3^flox^ breeder mice were purchased from the Jackson Laboratory. Prior to breeding Aldh1l1-Cre/ERT2 to Stat3^flox^, they were backcrossed 3 generations to C57BL/6J. Experimental mice were hemizygous Aldh1l1-Cre/ERT2; homozygous Stat3^flox^ (Cre +) and control mice were homozygous Stat3^flox^ (Cre-). All mice were treated i.p. with 75 mg/kg tamoxifen (MilliporeSigma) in corn oil once daily for 5 days at 7 weeks of age to induce Cre/ERT2 activity. Experiments were performed on male and female mice aged 12-24 weeks old. Mice were grouped-housed in temperature-and humidity-controlled rooms with free access to food and water under a 12-h light/dark cycle, unless they were undergoing ethanol or sucrose consumption tests, in which case they were individually housed on a reverse light/dark cycle. Experimental rooms were maintained at an ambient temperature of 21 ± 1°C with 40-60% humidity. All procedures were approved by the Institutional Animal Care and Use Committees of Virginia Commonwealth University and The Scripps Research Institute and complied with the ARRIVE guidelines and the National Institutes of Health *Guide for the Care and Use of Laboratory Animals*.

### 2.2 Tissue collection for RNA and protein analysis

Mice were euthanized by rapid decapitation. Brains were removed from the cranium and either rapidly frozen on dry ice or cut into 1 mm-thick sections on ice using a stainless-steel brain matrix (Zivic Instruments Inc., Pittsburgh, PA). The medial PFC (containing anterior cingulate, infralimbic and prelimbic regions, ∼1.9-0.9 mm anterior to bregma) was dissected using a razor blade, transferred to 1.5 ml centrifuge tubes, snap frozen and stored at −80°C until processing.

### 2.3 BaseScope^TM^ in situ hybridization

Brains were cut into 10 μm sections using a cryostat, mounted onto Superfrost Plus slides (ThermoFisher Scientific) and stored at −80°C until processing. Sections were fixed in 4% paraformaldehyde for 15 min at 4°C and then dehydrated with increasing ethanol concentrations (50, 70, and 100%) for 5 min at room temperature. Sections were treated with hydrogen peroxide for 8-10 min at room temperature and then with RNAScope Protease III (Advanced Cell Diagnostics, Inc., Newark, CA USA). Probes to *Aldh1l1* (BA-Mm-Aldh1l1-3zz-st-C1, #1048771-C1) and *Stat3* (BA-Mm-Stat3-3zz-st1-C2, #1279951-C2) were hybridized, amplified and detected using the BaseScope Duplex Assay^TM^ kit (Advanced Cell Diagnostics) according to the manufacturer’s instructions. Slides were counterstained with Gill’s hematoxylin. Brightfield images were acquired at 40X magnification using an Axioscope A1 microscope (Zeiss) equipped with a CCD camera. *Aldh1l1* and *Stat3* positive cells were counted manually by an investigator blinded to genotype from 6 sections per mouse and 3 mice per sex and genotype. Data are presented as the sum of the number of cells from all 6 sections or the percentage of *Aldh1l1*+, *Stat3* + cells from each mouse.

### 2.4 Quantitative real-time PCR (qPCR)

Total RNA was extracted using QIAzol lysis reagent (Qiagen, Germantown, MD, USA) followed by purification using the RNeasy Mini Kit (Qiagen). cDNA was synthesized using the High-Capacity cDNA Reverse Transcription Kit (ThermoFisher), and qPCR was performed using the SsoAdvanced Universal SYBR Green Supermix (Bio-Rad Laboratories, Hercules, CA, USA) on a CFX Duet Real-Time PCR System (Bio-Rad Laboratories). Primer sequences can be found in **Supplementary Table 1.** Relative expression was calculated using the 2^−ΔΔCt^ method by first subtracting the geometric mean of the Ct of three reference genes (*Gusb*, *Hprt*, and *Rpl13a*) from the Ct of the sample to obtain the ΔCt. The ΔΔCt was calculated by subtracting the average ΔCt of the male Cre - mice from the ΔCt of each sample.

### 2.5 Western blot

Frozen tissue was lysed and homogenized using a TissueLyser in ice cold 1X RIPA buffer (Cell Signaling Technology, Danvers, MA USA) with 1X Halt protease and phosphatase inhibitors (ThermoFisher Scientific, Waltham, MA). Protein concentrations were determined using the Pierce BCA Protein Assay kit (Thermo Fisher Scientific). Equal amounts of protein (15 μg) in SDS loading buffer with β-mercaptoethanol were separated by gel electrophoresis on precast 4%–12% NuPAGE™ Bis-Tris Mini Protein gels (ThermoFisher Scientific) and transferred to nitrocellulose membranes using the Miniblot Module (ThermoFisher Scientific) Membranes were blocked with 5% BSA in TBST. Membranes were incubated with primary (4°C, overnight) and secondary (1 h, room temperature) antibodies in 5% BSA in TBST (20 mM Tris, 150 mM NaCl, pH 7.4, 0.1% Tween-20). Primary antibodies were the following: EAAT2/GLT-1 (ERP5K) rabbit monoclonal (#20848S, Cell Signaling Technology), β-actin mouse monoclonal (#A5441, MilliporeSigma). Secondary antibodies were the following: donkey anti-mouse IgG (H+L) DyLight™ 680 (#SA510170, Invitrogen), IRDye® 800CW goat anti-rabbit IgG (#92632211, LI-COR). Blots were imaged on an Azure 500 Near-Infrared Fluorescent Imager (Azure Biosystems, Dublin, CA USA). Band intensities were analyzed using ImageJ (National Institutes of Health).

### 2.6 Electrophysiology

Electrophysiology studies were conducted in male 14-18 week old Stat3 aKO (Cre +, n=4) and controls (Cre-, n=4). Preparation of acute brain slices and electrophysiological recordings were performed as described previously (Patel RR, 2024; Patel et al., 2021; Patel et al., 2022; Roberts et al., 2019; Varodayan et al., 2023). Coronal slices (300 μm) of the prelimbic PFC were sectioned (Leica VT1200 S; Buffalo Grove, IL) from anesthetized mice in an ice-cold, high sucrose cutting solution (in mM): 206 sucrose; 2.5 KCl; 0.5 CaCl_2_; 7.0 MgCl_2_; 1.2 NaH_2_PO_4_; 26 NaHCO_3_; 5 HEPES; and 5 glucose). Slices recovered in artificial cerebrospinal fluid (aCSF; in mM): 130 NaCl, 3.5 KCl; 1.25 NaH_2_PO_4_; 1.5 MgSO_4_·7H_2_O; 2.0 CaCl_2_; 26 NaHCO_3_; and 10 glucose equilibrated with 95%O_2_/5% CO_2_ at 32 °C for 30 minutes followed by an incubation at RT for a minimum of 30 minutes before recording.

The slices were superfused (flow rate ∼2 mL/min) with aCSF, and all recordings were performed at room temperature (25±1.0 °C). Prelimbic pyramidal neurons (layer 2-3) were identified by their characteristic size and shape using infrared-differential interference contrast (IR-DIC) optics (Olympus BX50WI or BX51WI) equipped with a CCD camera (EXi Aqua, QImaging or equivalent), and only neurons with capacitance C_m_≥70 pF were used (as determined by initial membrane test (Clampex 11.2. acquisition software). The neurons were voltage clamped at-70 mV, and we used patch pipettes (3-6 MΩ) pulled from borosilicate glass (model no. G85150T-4, Warner Instruments, Hamden, CT, USA) and filled with K-gluconate internal solution (in mM): 145 K-gluconate, 0.5 EGTA, 2 MgCl_2_ 10 HEPES, 2.0 Mg-ATP, 0.2 Na-GTP. Before recording spontaneous synaptic currents, we determined the resting membrane potential (Vm) in current clamp mode, followed by current-voltage (I-V) tests to characterize basic intrinsic membrane properties and spiking characteristics. The I-V protocol comprised of 43 600 ms hyperpolarizing and depolarizing steps (10 pA steps), starting from-120 pA (Patel et al., 2021; Patel et al., 2022; Varodayan et al., 2023; Vlkolinsky, Khom, Vozella, Bajo, & Roberto, 2024). Whole-cell patch-clamp recordings of spontaneous excitatory postsynaptic currents (sEPSCs) were recorded in the presence of the GABA receptor antagonists 1 μM CGP 55845A (Tocris Biosciences, Ellisville, MI) and 10 μM picrotoxin (Ptx; MilliporeSigma, St. Louis, MO) dissolved in DMSO. Gap-free recordings of synaptic currents typically lasted 3-5 min and access (R_a_) resistance was continuously monitored with frequent 10-mV pulses. Neurons exhibiting changes of R_a_ >20% or with R_a_ > 20 MΩ were excluded from analysis.

### 2.7 Electrophysiological data analysis

To analyze synaptic currents, we measured frequencies, amplitudes, and rise and decay times, and areas of sEPSCs. Each synaptic event was visually confirmed and analyzed using MiniAnalyses Program v.6.0.8 (BlueCell, Korea). We averaged sEPSC characteristics in 3 min bins.

Intrinsic membrane properties (**Table 1**) and spike properties (I-V tests, **Table 2**) were analyzed using NeuroExpress 22.5.04 software (developed and kindly provided by Dr. A. Szucs) (Khom et al., 2020; Szucs & Huerta, 2014; Vlkolinsky et al., 2024). Briefly, at each current step, we measured the membrane potential (V_m_) difference and calculated passive membrane properties, such as membrane resistance (R_m_), capacitance (C_m_), time constant (τ_m_), membrane voltage sag (V_m_sag), and afterdepolarization (ADP_m_) by standard methods. We constructed linear fits of these values for each individual neuron and extrapolated them at *I* = 0 pA. We used these extrapolated values (reported as R_m_0, C_m_0, τ_m_0, V_sag_0, and ADP_m_0; for detailed description of calculation methods see (Vlkolinsky et al., 2024)) to calculate group averages and statistical significance for the Cre - and Cre + groups. If a large synaptic current incidentally interfered with the determination of an individual membrane parameter (*e.g.*, C_m_), then we excluded that neuron from analyses of the affected parameter only, but it remained included in analyses of other membrane parameters. Therefore, in I-V tests, the sample size varied for different intrinsic membrane parameters per group.

**Table 1.** Effect of genotype on intrinsic membrane properties in layer 2/3 prelimbic PFC pyramidal neurons of Cre – (control) and Cre + (Stat3 aKO) mice. Recordings were performed with KCl - based internal solution. Between group comparisons by two-tailed t-test, *p<0.05. The data are expressed as the mean ± SEM.

|  | **Cre – (n=47)** | **Cre + (n=62)** | **Statistics** |
| --- | --- | --- | --- |
| **Vrest (membrane resting potential, mV)** | -71.02 ± 0.51 | -70.82 ± 0.33 | t=0.3439, df=107,<br>p=0.732 |
| **Rmax (membrane resistance, MΩ)** | 219.5 ±<br>11.24 | 232.6 ± 9.63 | t=0.8831, df=107<br>p = 0.379 |
| **Time constant (membrane time constant, ms)** | 32.05 ± 1.61 | 33.23 ± 1.56 | t=0.5189, df=102<br>p = 0.61 |
| **Capacitance (membrane capacitance, pF)** | 150.7 ± 6.52 | 149.0 ± 4.47 | t=0.2277, df=104<br>p = 0.820 |
| **Membrane voltage sag slope (mV/nA)** | **-4.01 ± 0.55** | **-6.37 ± 0.91** | **t=2.029, df=106**<br>**\*p = 0.045** |
| **Afterdepolarization slope (mV/nA)** | -2.625 ± 1.50 | -4.92 ± 1.37 | t=1.125, df=101<br>p = 0.263 |

**Table 2.** Effect of genotype on AP characteristics in layer 2/3 prelimbic PFC pyramidal neurons of Cre – (control) and Cre + (Stat3 aKO) mice. Between group comparisons by two-tailed t-test, *p<0.05. The data are expressed as the mean ± SEM.

|  | **Cre – (n=47)** | **Cre + (n=62)** | **Statistics** |
| --- | --- | --- | --- |
| **Rheobase (pA)** | 76.99 ± 4.43 | 80.56 ± 4.90 | t=0.52, df=105<br>p = 0.605 |
| **Threshold (mV)** | -38.89 ± 0.36 | -38.30 ± 0.29 | t=1.28, df=103<br>p = 0.204 |
| **Amplitude (mV)** | 85.81 ± 1.06 | 85.14 ± 0.96 | t=0.47, df=102<br>p = 0.640 |
| **Half-width (mV)** | 1.59 ± 0.09 | 1.51 ± 0.06 | t=0.73, df=107<br>p = 0.468 |
| **Kink** | 291.6 ± 25.25 | 359.9 ± 27.48 | t=1.78, df=107<br>p = 0.081 |
| **Kink slope** | **4.42 ± 0.34** | **5.70 ± 0.36** | **t=2.52, df=107**<br>**\*p = 0.014** |
| **Upshoot slope (Vm/ms)** | 246.5 ± 12.15 | 251.5 ± 10.03 | t=0.32, df=107<br>p = 0.747 |
| **Spike positive phase** | 0.530 ± 0.01 | 0.518 ± 0.01 | t=0.93, df=107<br>p = 0.354 |
| **Decay slope (Vm/ms)** | -53.76 ± 2.69 | -53.91 ± 2.08 | t=0.04, df=107<br>p = 0.965 |
| **Spike negative**<br>**phase** | $0.801 \pm 0.01$ | $0.808 \pm 0.01$ | $t=0.61, df=107$<br>$p = 0.546$ |

### 2.8 Open-field locomotor activity and light-dark box tests

Following a 1 hour acclimation to the behavioral room in their home cages, mice were placed in individual open field chambers (26.7 × 26.7 × 20.3 cm Plexiglas boxes). Chambers were connected to a computerized three-dimensional activity monitoring system (Fusion v5.3; Omnitech Electronics Inc.) that recorded infrared photobeam breaks. Locomotor activity was measured for 30 minutes. Mice were tested two days later in the light-dark box. The open-field Plexiglas boxes were divided into equally sized light or dark compartments (25 × 12 × 20 cm each) using a black plastic partition with an opening in the middle for light–dark transitions. Preference for the light or dark side was assessed by measuring the time mice spent in each chamber in a 5 minute test session.

### 2.9 Ethanol and sucrose consumption tests

Following locomotor and light-dark box tests, mice were transferred to a reverse dark/light cycle room (lights off at 10 am and on at 10 pm) and acclimated for two weeks to the reverse dark/light cycle. They then underwent a modified ethanol drinking in the dark (DID) (Hamada et al., 2021) procedure in their home cages for three weeks. Three days prior to starting DID and throughout the test, mice were individually housed. Each mouse was given access to a single 10 ml sipper tube containing 15% ethanol (v/v), 3 hours into the dark cycle, for 4 hours on Mon-Thurs. The volume of fluid consumed was measured at the end of each session. On Fri-Sun, mice were not given ethanol. DID sessions resumed on Mon for a total of 3 weeks and mice were weighed weekly on Mon. To test sucrose consumption, a separate cohort of mice underwent four days of DID (Mon-Thurs), except the sipper tubes contained 2% sucrose in water instead of ethanol.

### 2.10 Ethanol loss-of-righting reflex (LORR)

Ethanol naïve mice were injected i.p. with 20% ethanol in saline (v/v) at a dose of 3.6 g/kg. Each mouse was placed on its back and tested for the ability to right itself. The mouse was determined to have lost the righting reflex if it could not right itself 3 times within 30 seconds and regained the righting reflex if it could fully right itself 3 times within 30 seconds. The duration of LORR was calculated as the difference between the time when the reflex was lost and when it was regained. One week later, mice were tested again for LORR at a dose of 4 g/kg ethanol.

### 2.11 Blood ethanol clearance assay

One week after testing LORR, mice were injected i.p. with 2 g/kg ethanol (20% v/v in saline). Five minutes after injection and then every 30 minutes after injection for a total of 3 hours, a 20 μl blood sample was taken from a tail vein puncture using a heparinized capillary tube. Blood was transferred into a 1.5 ml centrifuge tube, snap frozen and stored at −80°C until processing for blood ethanol concentrations (BECs). BECs were measured using the nicotinamide adenine dinucleotide-alcohol dehydrogenase (NAD-ADH) enzymatic assay as previously described (Zapata, Gonzales, & Shippenberg, 2006). β-NAD and yeast ADH were purchased from MilliporeSigma (St. Louis, MO, USA).

### 2.12 Statistical analysis

Unless otherwise indicated, data are presented as the mean ± SEM. All statistical analyses were performed using Prism software (version 10.1.1 or higher, GraphPad, San Diego, CA). For qPCR, locomotor activity, and light-dark box testing, a two-way ANOVA was used with between subject factors of genotype and sex, followed by post-hoc Sidak’s multiple comparisons tests when a significant interaction was observed. For Western blots and electrophysiology, data were analyzed by two-tailed *t-*test with Welch’s correction. Ethanol, sucrose and BEC data were analyzed by two-way repeated measures ANOVA separately for each sex with factors of time (within subject) and genotype (between subject). Mixed effects analysis was performed for male ethanol and sucrose consumption data because of one missing data point due to bottle leakage for one mouse on one day. Full statistical test results are reported in **Supplementary Table 2**.

## 3. Results

### 3.1 Stat3 deletion in adult astrocytes

To delete *Stat3* in astrocytes (Stat3 aKO), we crossed the tamoxifen-inducible Cre line, Aldh1l1-Cre/ERT2, with Stat3^flox^ mice and generated hemizygous Aldh1l1-Cre/ERT2; homozygous Stat3^flox^ (Cre +) mice. We treated 7-week-old mice with tamoxifen to activate Cre/ERT2 and induce recombination at loxP sites at the *Stat3* locus. Control mice were homozygous Stat3^flox^ (Cre-) mice treated with tamoxifen at the same age. To verify deletion of the floxed *Stat3* exons in astrocytes, we performed the BaseScope^TM^ Duplex assay in frontal cortex sections with specific probes to the floxed *Stat3* exons and *Aldh1l1* at least three weeks after tamoxifen injections (**Fig. 1A**). The number of *Aldh1l1*-expressing cells did not differ between genotypes (**Fig. 1B, t**_(7.205)_ = 0.469, *p* = 0.653), but the percentage of *Aldh1l1* + cells expressing *Stat3* was significantly reduced in Cre + mice (**Fig. 1C, t**_(9.885)_ = 4.422, *p* = 0.0013). The number of *Stat3*-expressing *Aldh1l1*-negative cells did not differ between genotypes (**Fig. 1D, t**_(9.999)_ = 1.445, *p* = 0.179). These results indicate selective deletion of *Stat3* in astrocytes in the cortex, as expected (Srinivasan et al., 2016). We noted that *Aldh1l1* is expressed in other organs, with high levels in the liver. We therefore measured *Stat3* expression by qPCR in the liver and found that *Stat3* expression was also significantly reduced in the liver of Cre + mice (**Supplementary Fig. 1**).

**Figure 1.**
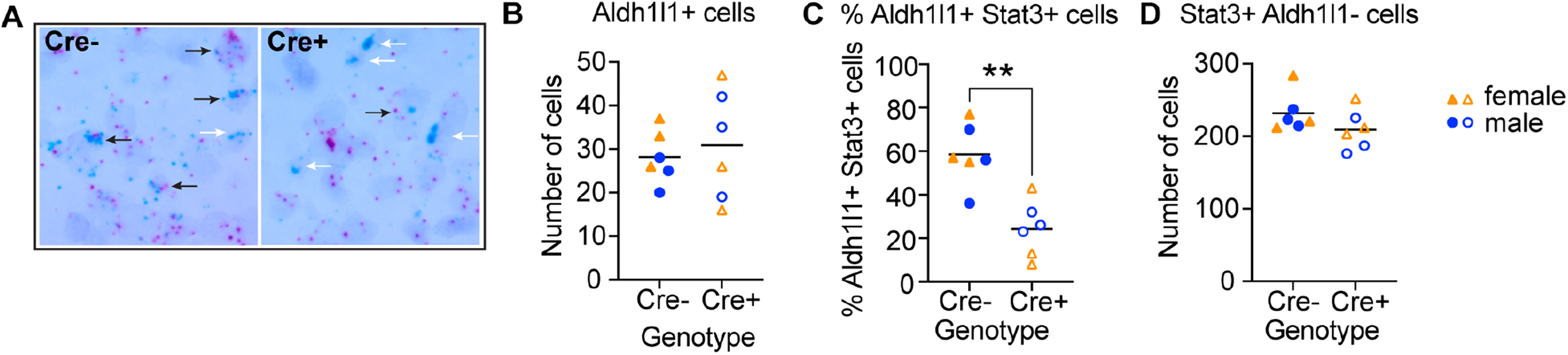
*Stat3* deletion in astrocytes. (**A**) Brightfield image of mouse cortex following BaseScope^TM^ Duplex Assay with probes to *Aldh1l1* (green) and *Stat3* floxed exons (pink) in Stat3 aKO (Cre +) and control (Cre-) mice. Black arrows point to *Aldh1l1-*expressing cells expressing *Stat3* floxed exons and white arrows point to *Aldh1l1*-expressing cells negative for *Stat3* floxed exons. (**B**) The number of *Aldh1l1*+ cells did not differ between genotypes. (**C**) The percentage of *Aldh1l1-*expressing cells expressing *Stat3* floxed exons was reduced in Cre + mice by student’s t-test, \*\**p*<0.01. (**D**) The number of *Stat3* floxed exon-expressing, *Aldh1l1*-negative cells did not differ between genotypes. For panels **B-D**, each point represents an individual mouse, with males in blue circles and females in orange triangles, n=3 mice per sex per genotype. The black line represents the mean.

### 3.2 Decreased Gfap and glutamate transporter transcripts in the PFC of Stat3 aKO mice

STAT3 regulates expression of genes encoding the astrocytic intermediate filament protein GFAP and astrocyte-enriched glutamate transporters (Bonni et al., 1997; Cheng et al., 2020; Feng et al., 2015; Ito et al., 2018; Moidunny et al., 2016; Shi et al., 2021; Wang et al., 2016). To determine if expression of these genes is affected by *Stat3* deletion in astrocytes, we used qPCR to measure mRNA levels in the PFC of Stat3 aKO and control mice (**Fig. 2**). *Gfap* expression was significantly reduced in both male and female Stat3 aKO compared with control mice (**Fig. 2A**, genotype, F_(1, 43)_ = 68.28, *p* < 0.0001; sex, F_(1, 43)_ = 2.381, *p* = 0.130; genotype × sex interaction, F_(1, 43)_ = 0.006, *p* = 0.941). We next examined the expression of the glutamate transporter genes *Slc1a2*, *Slc1a3*, *Slc7a11* and *Slc17a8*. *Slc1a2* mRNA was significantly reduced in male and female Stat3 aKO compared with control mice (**Fig. 2B**, genotype, F_(1, 45)_ = 7.800, *p* = 0.0076; sex, F_(1, 45)_ = 0.054, *p* = 0.818; genotype × sex interaction, F_(1,45)_ = 0.322, *p* = 0.573). *Slc1a3* expression was slightly reduced in Stat3 aKO mice, but the reduction was not statistically significant (**Fig. 2C**, genotype, F_(1, 44)_ = 3.370, *p* = 0.073; sex, F_(1, 44)_ = 0.154, *p* = 0.697; genotype × sex interaction, F_(1, 44)_ = 0.423, *p* = 0.519). *Slc7a11* expression exhibited a significant genotype × sex interaction, with expression slightly decreased in male and increased in female Stat3 aKO mice (**Fig. 2D**, genotype, F_(1, 44)_ = 0.437, *p* = 0.512; sex, F_(1, 44)_ = 0.021, *p* = 0.886; genotype × sex interaction, F_(1, 44)_ = 4.891, *p* = 0.032). However, post hoc multiple comparisons testing did not reveal significant differences between Stat3 aKO and controls in either males (*p* = 0.453) or females (*p* = 0.110). Finally, *Slc17a8* expression was reduced in Stat3 aKO compared with control mice (**Fig. 2E**, genotype, F_(1,44)_ = 7.111, *p* = 0.011; sex, F_(1,44)_ = 0.020, *p* = 0.889; genotype × sex interaction, F_(1,44)_ = 2.209, *p* = 0.144). Given that *Stat3* deletion in astrocytes resulted in decreased expression of *Slc1a2* and *Slc17a8*, we hypothesized that STAT3 might directly regulate these genes through STAT3 binding motifs in their promoters. We uncovered 4 strongly predicted STAT3 binding motifs within the 2 kb upstream promoter through the first intron of *Slc1a2* (**Supplementary Table 3**). Together, these results indicate that STAT3 in PFC astrocytes regulates the expression of *Slc1a2*, *Slc17a8* and *Gfap*, possibly through direct transcriptional regulation.

**Figure 2.**
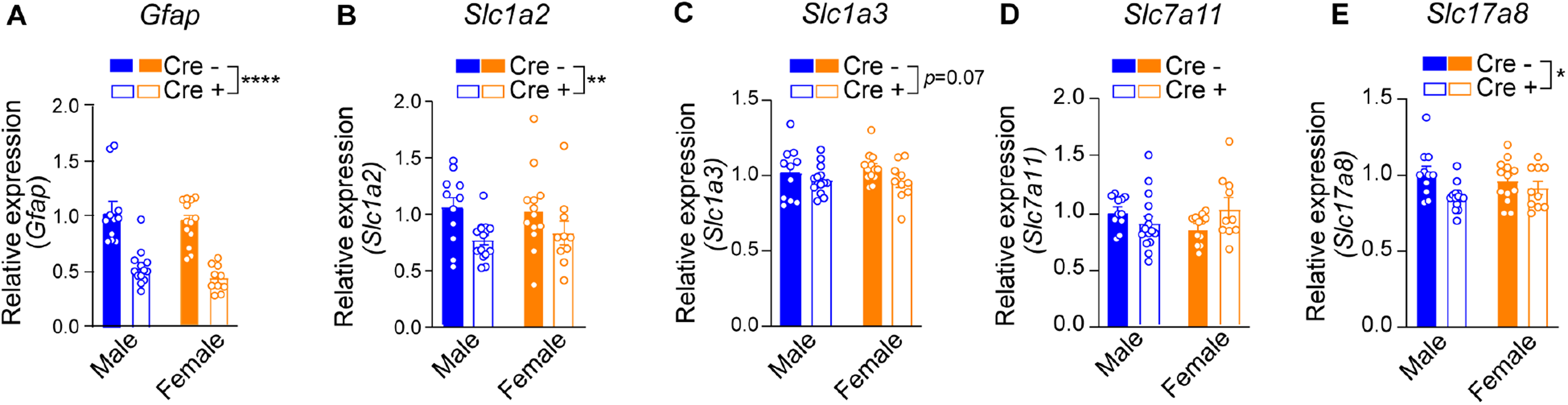
Decreased *Gfap* and glutamate transporter transcripts in the PFC of Stat3 aKO mice. RNA was extracted from the PFC and qPCR was performed on cDNA from Stat3 aKO (Cre +) and control (Cre-) mice. Relative expression of (**A**) *Gfap*, (**B**) *Slc1a2*, (**C**) *Slc1a3*, (**D**) *Slc7a11*, and (**E**) *Slc17a8*. Data are shown as the mean ± SEM (n = 9–11 per group). *p < 0.05, **p < 0.01, ****p < 0.0001, main genotype effect by two-way ANOVA.

### 3.3 Decreased GLT-1 protein in the PFC of male Stat3 aKO mice

We next focused on *Slc1a2*, as this gene encodes the astrocyte-enriched glutamate transporter GLT-1/EAAT2, which is responsible for 95% of glutamate uptake activity in the young adult forebrain (Danbolt, Storm-Mathisen, & Kanner, 1992; Otis & Kavanaugh, 2000; Tanaka et al., 1997) and showed the greatest reduction in transcript levels in Stat3 aKO mice. To assess whether the decrease in *Slc1a2* mRNA in the PFC of Stat3 aKO mice was reflected by reduced protein, we performed western blotting for GLT-1 in PFC homogenates from Stat3 aKO and control mice (**Fig. 3A & D**). We observed both GLT-1 monomeric (∼65 kDa) and dimeric forms (∼130 kDa) on the blots, as previously reported (Haugeto et al., 1996). GLT-1 monomers did not differ between genotypes in either males or females (**Fig. 3B & E**: males, t_(6.55)_ = 0.458, *p* = 0.662; females, t_(4.24)_ = 0.828, *p* = 0.452). In males, GLT-1 dimers were significantly decreased in Stat3 aKO mice compared with controls (**Fig. 3C, t**_(7.18)_ = 3.204, *p* = 0.014). Interestingly, in females, GLT-1 dimers were not altered by *Stat3* deletion in astrocytes (**Fig. 3F, t**_(6.81)_ = 0.775, p = 0.465). These results indicate that *Stat3* deletion in astrocytes reduced GLT-1 protein in the PFC of males but had no effect in females.

**Figure 3.**
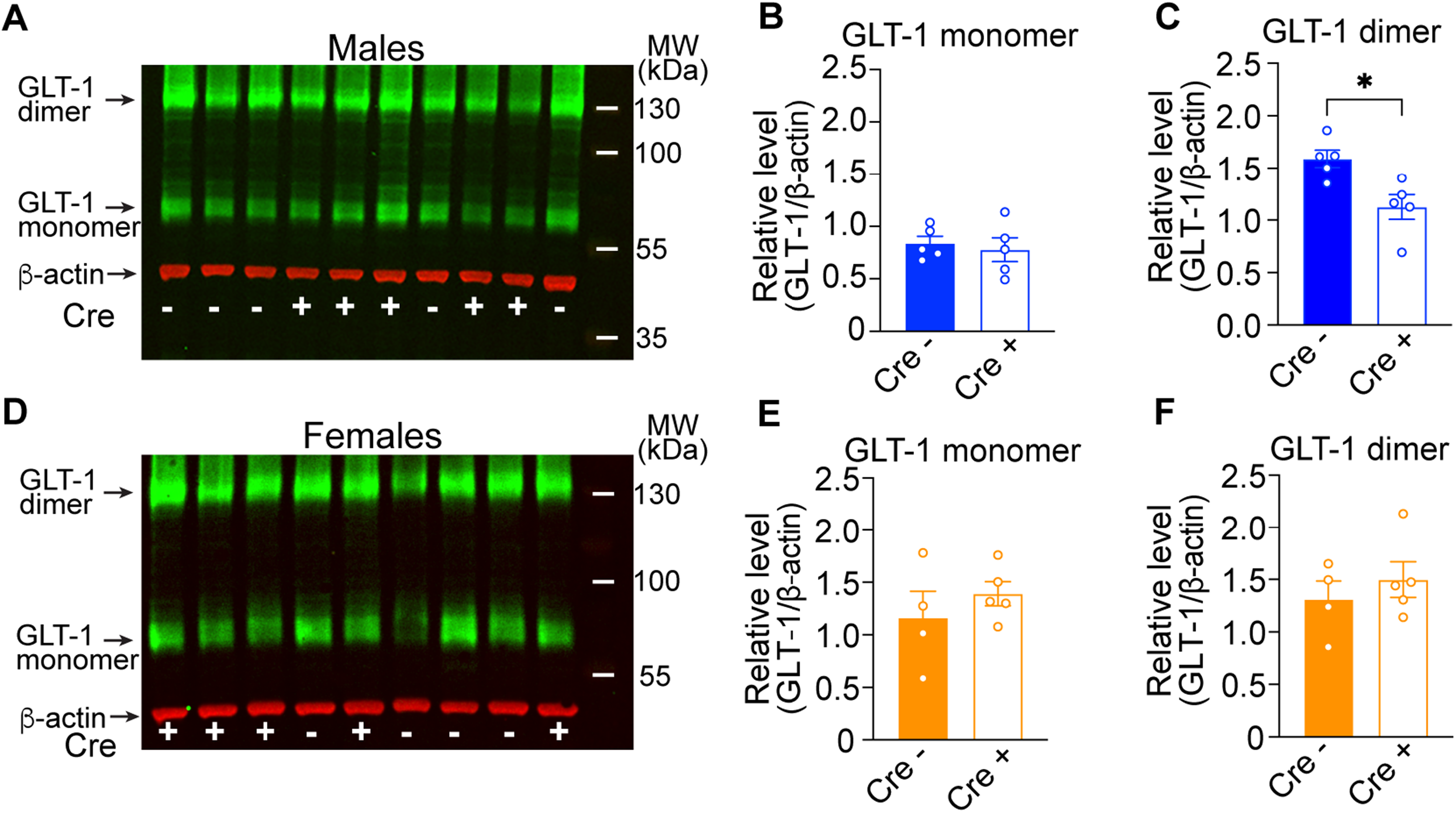
Decreased GLT-1 protein in the PFC of male Stat3 aKO mice. Protein lysates from the PFC of Stat3 aKO (Cre +) and control (Cre-) mice were immunoblotted with antibody to GLT-1. (**A, D**) Representative fluorescent multiplex western blots of GLT-1 in (**A**) male and (**D**) female mice. Arrows point to green GLT-1 dimer (∼130 kDa) and monomer (∼65 kDa), and red β-actin (∼42 kDa) bands. Cre genotype (-or +) is labeled at the bottom of the blot and molecular weight (MW) markers are labeled with white bars on the right of the blot. (**B–C**) GLT-1 dimer and monomer quantification in males, normalized to β-actin. (**E–F**) Same in females. Data are shown as the mean ± SEM (n = 4–5 per group). \**p* < 0.05, two tailed *t*-test with Welch’s correction.

### 3.4 Enhanced baseline spontaneous glutamatergic neurotransmission in prelimbic pyramidal neurons of Stat3 aKO mice

Given the sex-specific effect of *Stat3* deletion in astrocytes on GLT-1 protein, we next determined if glutamatergic neurotransmission is altered in male Stat3 aKO mice. We hypothesized that Stat3 aKO mice would exhibit enhanced glutamate transmission resulting from decreased glutamate clearance. We recorded glutamate receptor-mediated sEPSCs in prelimbic pyramidal neurons in male control and Stat3 aKO mice (**Fig. 4**). We found a significant increase in the mean baseline sEPSC amplitude (**Fig. 4C, t**_(51)_ = 2.836, *p* = 0.0065) in Stat3 aKO compared with control mice. There were no differences in the baseline frequency (**Fig. 4B**), rise and decay kinetics (**Fig. 4D, E**), or area (**Fig. 4F**) of sEPSCs between genotypes. Intrinsic passive and active membrane properties of the prelimbic pyramidal neurons did not differ significantly between Stat3 aKO and control mice on all measures (**Table 1**). These results indicate that deletion of *Stat3* in adult male astrocytes, associated with reduced GLT-1 levels, results in potentiated glutamatergic synaptic transmission in the prelimbic cortex, likely due to a reduction in astrocyte glutamate uptake activity.

**Figure 4.**
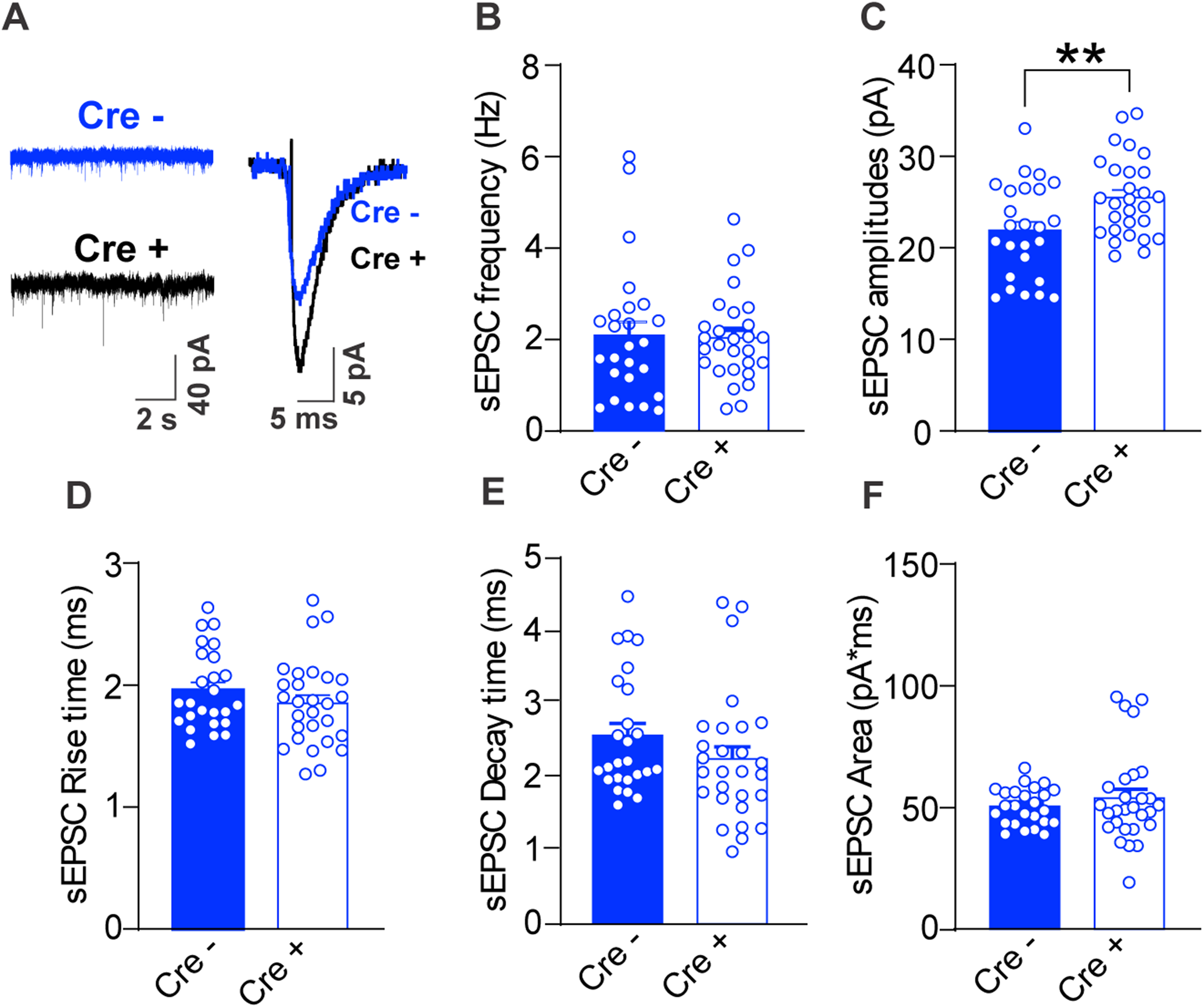
Enhanced baseline spontaneous glutamatergic transmission in prelimbic pyramidal neurons of male Stat3 aKO mice. (**A**) Representative traces of sEPSCs from a prelimbic cortex neuron (left panel) and average amplitude of sEPSCs (right panel) of control (Cre-) and Stat3 aKO (Cre +) mice. (**B**) Baseline frequency did not differ between genotypes. (**C**) Amplitude was significantly increased in Cre + compared with Cre - mice, \*\**p*=0.0065, t(51) = 2.836 by unpaired two-tailed t-test. (**D**) Rise time, (**E**) decay time, and (**F**) area of sEPSCs did not differ between genotypes. Cre-, n= 25 cells from 4 mice and Cre +, n = 28 cells from 4 mice. Data are plotted as the mean ± SEM.

### 3.5 Baseline spontaneous GABAergic transmission and intrinsic membrane properties of central nucleus of the amygdala (CeA) neurons are not altered in Stat3 aKO mice

We next assessed whether *Stat3* deficiency in astrocytes was associated with alterations in GABA_A_ receptor-mediated synaptic transmission in the CeA, a brain region enriched in GABAergic inhibitory neurons. Note that CeA slices were obtained from the same male Stat3 aKO and control mice that were used for the sEPSC recordings in the prelimbic cortex slices. In CeA, we recorded action-potential dependent, spontaneous inhibitory post-synaptic currents (sIPSCs) and membrane properties. We did not find any effect of genotype on sIPSC frequency, amplitude, rise, decay or area (**Supplementary Figure 2B-F),** indicating that deletion of *Stat3* from adult male astrocytes does not alter action-potential-dependent GABA release in the CeA. Intrinsic passive and active membrane properties of the CeA neurons did not differ significantly on all measures calculated between the Stat3 aKO and control mice (**Supplementary Tables 4 & 5**).

### 3.6 Stat3 deletion in astrocytes does not affect baseline locomotor or anxiety-like behaviors

To assess whether *Stat3* knockout in astrocytes has behavioral consequences, we measured baseline locomotor and anxiety-like behavior in Stat3 aKO mice. Mice were placed in a locomotor activity chamber for 30 minutes. The distance traveled, time spent moving and time in the center of the chamber did not differ by genotype or sex (**Supplementary Fig. 3A-C**). We tested the mice in a light-dark box two days later. Stat3 aKO mice did not differ in time spent on the light side of the box compared with control mice (**Supplementary Fig. 3D**). These results indicate that Stat3 aKO mice do not have a gross locomotor or anxiety-like behavioral phenotypes.

### 3.7 Male Stat3 aKO mice consume less ethanol than controls

We previously found that mice treated with a STAT3 inhibitor consume less ethanol than vehicle-treated mice in the DID binge drinking test (Hamada et al., 2021), and other groups have found that GLT-1 activity is associated with ethanol consumption in mice and rats (Lee et al., 2013; Tan et al., 2026). To determine whether STAT3 in astrocytes regulates ethanol consumption, Stat3 aKO and controls of both sexes underwent the drinking in the dark (DID) test for binge ethanol consumption for 12 drinking sessions over three weeks. Male Stat3 aKO mice consumed less ethanol than control males (**Fig. 5A**: genotype, F_(1, 17)_ = 10.95, *p* = 0.0041; time, F_(11, 186)_ = 18.04, *p* <0.0001; genotype x time interaction, F_(11, 186)_ = 0.5259, *p* = 0.8842), whereas ethanol intake by female Stat3 aKO mice was comparable to control females (**Fig. 5D**). Stat3 aKO mice body weights were not significantly different than sex-matched controls over the three weeks of ethanol consumption testing (**Supplementary Fig. 3G**). We next determined whether the effect of *Stat3* deficiency in astrocytes on ethanol consumption generalized to other fluids by testing 2% sucrose consumption over 4 days in the DID procedure. Neither male nor female Stat3 aKO mice exhibited significant differences in sucrose intake (**Fig. 5B, E**). We next tested a separate cohort of Stat3 aKO mice for ethanol-induced loss-of-righting reflex (LORR) at two ethanol doses (3.6 and 4 g/kg, i.p., spaced one week apart). Stat3 aKO mice did not differ from controls in LORR (**Supplementary Fig. 3E, F**). In the same mice that were tested for LORR, we tested blood ethanol clearance one week later with a 2 g/kg i.p. ethanol injection. There was no effect of genotype on blood ethanol clearance in either sex (**Fig. 5C, F**). Finally, a separate group of mice was tested for ethanol conditioned place preference (CPP) with a 2g/kg i.p. ethanol dose (see **Supplementary Methods**). Stat3 aKO mice did not differ in ethanol CPP when compared to their control counterparts (**Supplementary Fig. 3H**). Together, these results indicate a sex-specific effect of *Stat3* deletion in adult male astrocytes on ethanol consumption that is not due to altered ethanol metabolism.

**Figure 5.**
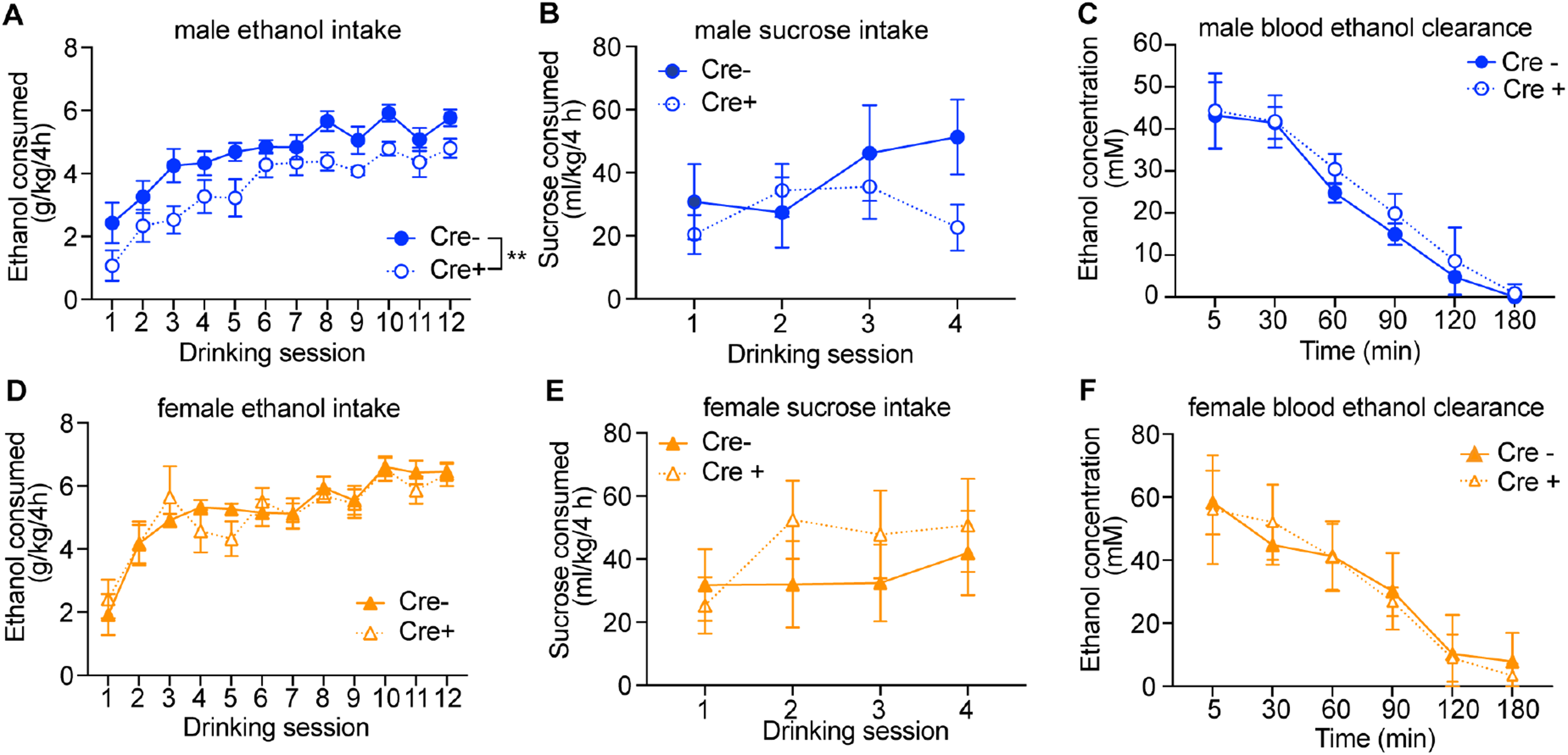
**Male Stat3 aKO mice consume less ethanol than control males**. (**A**) Ethanol consumption in g/kg by male Stat3 aKO (Cre +, n =9) and control (Cre-, n=10) mice in the drinking in the dark test over 12 sessions. Cre + males consumed significantly less ethanol than Cre – males \*\**p*=0.0041, main effect of genotype by mixed-effects ANOVA. (**B**) Sucrose consumption by male mice (n=11 Cre - and 13 Cre +) over 4 drinking sessions. (**C**) Blood ethanol concentrations (in mM) in male Cre – (n = 5) and Cre + (n = 6) mice after i.p. injection of 2 g/kg ethanol. (**D**) Ethanol consumption in g/kg by female mice (n=8 Cre – and 11 Cre +) in the DID test over 12 sessions. (**E**) Sucrose consumption by female mice (n=8 Cre – and 11 Cre +) over 4 drinking sessions. (F) Blood ethanol concentrations (in mM) in female Cre – and Cre + mice (n = 6 per genotype) after i.p. injection of 2 g/kg ethanol. In panels **A, B, D, E**, data are plotted as the mean ± SEM; in panels **C & F**, data are plotted as mean ± SD.

## 4. Discussion

Here we have shown that STAT3 in astrocytes regulates the expression of *Gfap, Slc1a2* and *Slc17a8* in the PFC of adult mice in a non-pathological state. The decrease in *Slc1a2* transcript was associated with reduced GLT-1 protein and an increase in glutamatergic neurotransmission in the PFC of male mice. To our knowledge, this is the first demonstration that genetic deletion of *Stat3* from adult astrocytes alters glutamate transporter expression and excitatory synaptic transmission *in vivo* under basal conditions in a sex-specific manner. Male, but not female Stat3 aKO mice consumed less ethanol than controls in a binge drinking test. These results demonstrate that astrocytic STAT3 is important for maintaining both cortical GLT-1 levels and driving excessive ethanol consumption in male mice.

*Stat3* knockout in astrocytes resulted in a reduction in *Slc1a2* mRNA; we found several predicted STAT3 binding motifs in the *Slc1a2* gene, suggesting that STAT3 promotes *Slc1a2* expression via direct DNA binding, like in *Gfap,* a known STAT3 target gene (Bonni et al., 1997; Ito et al., 2018). An open question is how STAT3 maintains the expression of these genes in astrocytes. The canonical model of transcriptional regulation by STAT3 is that it is phosphorylated at tyrosine 705 (pySTAT3) upon growth factor and cytokine stimulation. Under basal conditions, pySTAT3 is typically very low or even undetectable, although it’s possible that a certain level of pySTAT3 is sustained by ongoing growth factor stimulation of astrocytes in the adult brain to maintain gene expression. Alternatively, unphosphorylated and/or lysine acetylated STAT3 may regulate gene expression in astrocytes under unstimulated conditions, as unphosphorylated STAT3 is also present in the nucleus and has been shown to regulate gene expression and chromatin structure (Dasgupta et al., 2014; Timofeeva et al., 2012).

Depletion of STAT3 in astrocytes resulted in reduced *Slc1a2* transcript in both sexes, yet GLT-1 protein was only decreased in male Stat3 aKO mice. Post-transcriptional mechanisms, such as increased protein translation, stability or trafficking could normalize GLT-1 protein levels in female Stat3 aKO mice to compensate for decreased mRNA levels. Surprisingly, only GLT-1 dimers, but not monomers, were decreased in male Stat3 aKO mice. This may have been due to a tissue processing artifact. The mature, active form of GLT-1 is a trimer (Haugeto et al., 1996; Kato et al., 2022), but this form is disrupted when fresh brain tissue is lysed with ionic detergents and reducing agents. Oxidation causes irreversible cross-linking of GLT-1, visualized as oligomers on gels (Haugeto et al., 1996). Our mouse brain samples were frozen prior to lysis and gel electrophoresis, so GLT-1 may have been cross-linked during tissue freezing and storage even though we used ionic detergents and reducing agents, and the decrease in GLT-1 dimers in Stat3 aKO males reflected an overall reduction in protein levels. We did not detect trimers or higher molecular weight oligomers on our western blots because the molecular weight upper limit of the gels was ∼260 kDa. Another possibility is that the decrease in GLT-1 dimers in male Stat3 aKO mice might indicate that STAT3, in addition to regulating gene transcription, is also involved in trafficking and assembly of GLT-1 at the plasma membrane. In primary cortical astrocytes, *Gfap* knockout resulted in decreased GLT-1 trafficking to the membrane (Hughes, Maguire, McMinn, Scholz, & Sutherland, 2004). The decrease in *Gfap* expression in male Stat3 aKO might impair GLT-1 trafficking, indicating that loss of STAT3 in astrocytes indirectly contributes to decreased GLT-1 dimer formation through a GFAP-related mechanism.

To ascertain the impact of decreased GLT-1 protein in male Stat3 aKO mice on glutamate neurotransmission, we measured sEPSCs in layer 2-3 pyramidal neurons in the prelimbic PFC. The sEPSC amplitude was increased in Stat3 aKO, consistent with decreased GLT-1 and impaired clearance of extracellular glutamate (Fullana, Covelo, Bortolozzi, Araque, & Artigas, 2019; Petr et al., 2015). Fullana et al found that siRNA-mediated knockdown of GLT-1 in the mouse infralimbic cortex resulted in increased sEPSC frequency and amplitude onto layer 5 pyramidal neurons (Fullana et al., 2019). These results are comparable to what we found in the prelimbic cortex of the Stat3 aKO, although sEPSC frequency was not changed in the Stat3 aKO. Other effects of STAT3 knockout in astrocytes may minimize or compensate for the reduction in GLT-1; for example, the decrease in *Slc17a8*, encoding VGLUT3, in Stat3 aKO, which might reduce calcium-dependent glutamate release from astrocytes. It is important to note that the effect of depleting STAT3 in astrocytes selectively impacted glutamatergic neurotransmission, as GABA_A_-mediated synaptic transmission in the CeA was unaffected in Stat3 aKO, although a caveat is that we did not measure GABAergic neurotransmission in the PFC.

STAT3 in astrocytes regulates levels of GLT-1 in different pathological states, but the direction of the effect differs from what we observed here under basal conditions. For example, pySTAT3 was increased in mouse hippocampal astrocytes during methamphetamine withdrawal, and this was associated with decreased GLT-1 protein. Viral-mediated knockdown of STAT3 in hippocampal astrocytes restored GLT-1 protein and normalized glutamate levels (Shi et al., 2021). Similarly, in primary cortical astrocytes, treatment with the cytokine oncostatin M (OSM) increased pySTAT3 and decreased GLT-1 expression and glutamate uptake; treatment with the JAK inhibitor AG 490, which decreases JAK-dependent STAT3 tyrosine phosphorylation, prevented the decrease in glutamate uptake (Moidunny et al., 2016). In contrast to these two studies which demonstrated that blocking or knocking down STAT3 restored GLT-1 levels, we found that knockout of STAT3 in astrocytes reduced *Slc1a2*/GLT-1 levels. These results underscore the importance of context when studying the function of STAT3. Under normal conditions, STAT3 is important for maintaining expression of *Slc1a2*, but during tissue injury or inflammation, activated STAT3 may instead undergo a transcriptional switch, resulting in repression of *Slc1a2* expression. Behavioral characterization of the Stat3 aKO mice indicated that baseline locomotor and anxiety-like behaviors were not altered in the Stat3 aKO, despite evidence that reduced GLT-1 in the PFC is associated with increased anxiety-like behavior (Khan, Ronan, & Rahman, 2026). We next tested the Stat3 aKO mice for binge ethanol consumption because we previously found decreased binge drinking in both male and female mice treated systemically with the STAT3 inhibitor stattic (Hamada et al., 2021). Male Stat3 aKO mice consumed less ethanol than male controls, indicating that astrocyte-expressed STAT3 promotes binge drinking in male, but not female mice. The lack of an effect on binge ethanol consumption in female Stat3 aKO mice contrasts with our findings with stattic, but i.p. administration of stattic inhibits STAT3 activity in all cells, not just astrocytes. The stattic results suggest that perhaps STAT3 functions in other cell types in females to regulate binge ethanol consumption.

The finding that male Stat3 aKO mice consume less ethanol raises two important questions. First, is there a specific region of the brain in which astrocyte-expressed STAT3 promotes ethanol intake? Several studies have shown that regionally manipulating astrocyte activity affects ethanol consumption (Erickson et al., 2021; Kastner-Blasczyk et al., 2025; Nwachukwu et al., 2021; Tan & Ding, 2026). Male PFC astrocytes have been shown to bidirectionally regulate ethanol intake (Erickson et al., 2021). Increasing calcium signaling in PFC astrocytes in ethanol-naïve male mice with a Gq-coupled designer receptor activated by designer drugs (DREADD) increased ethanol consumption, and decreasing calcium signaling by expressing a calcium extruder (plasma membrane calcium ATPase) in PFC astrocytes reduced the escalation of ethanol consumption (Erickson et al., 2021). Another region where astrocyte activity drives ethanol consumption is the orbitofrontal cortex (OFC) (Kastner-Blasczyk et al., 2025). Mice exposed to chronic intermittent ethanol vapor exposure escalate their ethanol intake; expression of a calcium extruder in OFC astrocytes prevented this escalation in male, but not female, mice, indicating a sex-specific effect of inhibiting astrocyte activity in the OFC on ethanol intake (Kastner-Blasczyk et al., 2025). Astrocytes in the basolateral amygdala (BLA) have also been implicated in binge ethanol consumption, as activation of BLA astrocytes by Gq-coupled DREADD reduced ethanol consumption in male mice (Nwachukwu et al., 2021). In the ventral striatum (nucleus accumbens) of male mice, expression of a calcium extruder in astrocytes increased aversion-resistant ethanol consumption, in which ethanol was adulterated with quinine (Peyton et al., 2025). Finally, STAT3 is increased in astrocytes in the CeA and nucleus accumbens of mice, and hippocampus of rats, after chronic ethanol exposure (Chen et al., 2021; Hashimoto et al., 2026; Hashimoto et al., 2025), implicating these regions as potential candidates for STAT3 activity in driving ethanol consumption. Future studies should identify the brain region(s) in which astrocytic STAT3 promotes binge ethanol drinking.

The second question raised by our finding that male Stat3 aKO mice consume less ethanol is whether the reduction in ethanol consumption is due to altered astrocyte GLT-1 expression and glutamate neurotransmission. Chronic ethanol exposure disrupts glutamate homeostasis, partly by downregulating the expression of GLT-1 in the mesocorticolimbic system (Alasmari, Goodwani, McCullumsmith, & Sari, 2018). The β-lactam compounds, ceftriaxone and MC-100093, restore GLT-1 levels and glutamate homeostasis after chronic ethanol drinking and are effective in reducing ethanol consumption in alcohol-preferring rats and mice (Alotaibi et al., 2024; Das, Yamamoto, Hristov, & Sari, 2015; Lee et al., 2013; Sari, Sreemantula, Lee, & Choi, 2013). Intra-ventricular infusion of the GLT-1 activator LDN-212320 into alcohol-preferring rats after chronic ethanol consumption also reduced ethanol consumption and preference (Tan et al., 2026). Based on the literature, we would have predicted that the Stat3 aKO mice would consume more ethanol, not less, given the decrease in GLT-1 levels and increased glutamate neurotransmission in the PFC. However, we do not know the effect of Stat3 aKO on GLT-1 levels in response to ethanol exposure, or in regions other than the PFC that are involved in binge ethanol consumption. More investigation is needed to sort out these questions. Together, these findings identify astrocytic STAT3 as a sex-specific regulator of glutamate transporter expression, excitatory synaptic transmission in the PFC, and voluntary ethanol intake. These results indicate an important role for astrocytic STAT3 *in vivo* in the maintenance of brain gene expression and behavior in the absence of inflammatory stimuli or injury.

## Data availability statement

The data that support the findings of this study are openly available in The Open Science Framework at https://doi.org/10.17605/OSF.IO/FDTVW

## Acknowledgements

This work was supported by NIH Grants U01AA020912, R01AA027231 and P50AA022537 to AWL; U01AA013498, R01AA029841 and P60AA006420 to MR; and T32AA007456 to CME. We would like to thank the VCU Massey Comprehensive Cancer Center Transgenic/Knockout Mouse Shared Resource for mouse breeding. The services were supported, in part, with funding from NIH-NCI Cancer Center Support Grant P30CA016059. We would also like to thank Bhavya Pulugujju for qPCR assistance.

