## Supplementary Material for "Astrocyte-expressed STAT3 regulates glutamate homeostasis and binge ethanol drinking in mice"

**Supplementary Methods**

*Electrophysiology of the central nucleus of the amygdala (CeA)*

Electrophysiology studies were conducted in male 14-18 weeks old *Stat3* astrocyte knockout (aKO) mice (n=4, 26.3 ± 2.2 g) and male wild-type littermates (n=4; 24.3 ± 1.0 g). Preparation of acute brain slices and electrophysiological recordings were performed as described previously (Patel RR, 2024; Patel et al., 2021; Patel et al., 2022; Roberts et al., 2019; Varodayan et al., 2023). Coronal slices (300 µm) containing central amygdala (CeA) were sectioned (Leica VT1200 S; Buffalo Grove, IL) from anesthetized mice in an ice-cold, high sucrose cutting solution (in mM): 206 sucrose; 2.5 KCl; 0.5 $CaCl_2$; 7.0 $MgCl_2$; 1.2 $NaH_2PO_4$; 26 $NaHCO_3$; 5 HEPES; and 5 glucose. Slices were superfused (flow rate ~2 mL/min) with artificial cerebrospinal fluid (aCSF; in mM): 130 NaCl, 3.5 KCl; 1.25 $NaH_2PO_4$; 1.5 $MgSO_4 \cdot 7H_2O$; 2.0 $CaCl_2$; 26 $NaHCO_3$; and 10 glucose equilibrated with 95%$O_2$/5% $CO_2$. Whole-cell patch-clamp recordings were performed in neurons from the medial subdivision of the central amygdala (CeA) voltage-clamped at −60 mV. All recordings were performed at room temperature (25±1.0 °C). Before recording spontaneous synaptic currents, we determined the resting membrane potential (Vm) in current clamp mode, followed by current-voltage (I-V) tests to characterize basic intrinsic membrane properties and spiking characteristics. The I-V protocol comprised of 43 600 ms hyperpolarizing and depolarizing steps (10 pA steps), starting from −120 pA (Khom, Steinkellner, Hnasko, & Roberto, 2020; Khom, Wolfe, et al., 2020; Roberts et al., 2019). I-Vs were conducted using the K-gluconate internal (in mM): 145 K-gluconate, 0.5 EGTA, 2 $MgCl_2$ 10 HEPES, 2.0 Mg-ATP, 0.2 Na-GTP. Glass patch pipettes (3-6 MΩ) were pulled from borosilicate glass (model no. G85150T-4, Warner Instruments, Hamden, CT, USA).

To record GABAergic spontaneous inhibitory post-synaptic currents (sIPSCs), we used patch pipettes filled with the KCl internal solution composed of (in mM): 135 KCl, 5.0 EGTA; 2

$MgCl_2$, 10 HEPES; 2 Mg-ATP; and 0.2 Na-GTP. sIPSCs were recorded in the presence of 6,7-dinitroquinoxaline-2,3-dione (DNQX, 20 μM), DL-2-amino-5-phosphonovalerate (DL-AP5, 30 μM), and CGP 55845A (1 μM). Gap-free recordings of synaptic currents typically lasted 3-5 min and access ($R_a$) resistance was continuously monitored with frequent 10-mV pulses. Neurons exhibiting changes of $R_a$ >20% or with $R_a$ > 20 MΩ were excluded from analysis.

*Electrophysiological data analysis*

To analyze synaptic currents, we measured frequencies, amplitudes, and rise and decay times, and areas of sIPSCs. Each synaptic event was visually confirmed and analyzed using MiniAnalyses Program v.6.0.8 (BlueCell, Korea). We averaged sIPSC characteristics in 3 min bins. Intrinsic membrane properties and spike properties (I-V tests, **Supplementary Tables 4 and 5**) were analyzed using NeuroExpress 22.5.04 software (developed and kindly provided by Dr. A. Szucs) (Khom, Wolfe, et al., 2020; Szucs & Huerta, 2014; Vlkolinsky, Khom, Vozella, Bajo, & Roberto, 2024). Briefly, at each current step, we measured the membrane potential ($V_m$) difference and calculated passive membrane properties, such as membrane resistance ($R_m$), capacitance ($C_m$), time constant ($\tau_m$), membrane voltage sag ($V_m$sag), and afterdepolarization ($ADP_m$) by standard methods. We constructed linear fits of these values for each individual neuron and extrapolate them at $I = 0$ pA. We used these extrapolated values (reported as $R_m0$, $C_m0$, $\tau_m0$, $V_{sag}0$, and $ADP_m0$; see (Vlkolinsky et al., 2024) for detailed descriptions of calculation methods) to calculate group averages and statistical significance for the Cre- control and Cre+ Stat3 aKO groups. If a large synaptic current incidentally interfered with the determination of an individual membrane parameter (*e.g.*, $C_m$), then we exclude that neuron from analyses of the affected parameter only, but it remained included in analyses of other membrane parameters.

Therefore, in I-V tests, the sample size varied for different intrinsic membrane parameters per group. All data are shown as mean ± SEM values and were statistically evaluated by GraphPad Prism version 10.1.2 or higher (GraphPad Software, San Diego, CA). Data were analyzed by two-tailed *t*-test.

*Ethanol conditioned place preference (CPP)*

CPP training and testing were conducted in a modified mouse open field apparatus with 48-channel infrared beam detectors and Activity Monitor software (Med Associates, St. Albans, VT). Modifications of the open field apparatus to create a 2-chambered setup for CPP have been described previously (Hilderbrand & Lasek, 2018). On the first day of testing, mice were placed in the CPP apparatus and allowed access to both sides of the chamber for 30 min to measure baseline preference. Mice were then assigned to the non-preferred side for ethanol injections. Mice were conditioned over the next 8 weekdays with an intraperitoneal (i.p.) injection of 2 g/kg ethanol (20% v/v in 0.9% saline) on days 2, 4, 6 & 8 or saline (10 ml/kg) on days 3, 5, 7 & 9. Immediately after injection with ethanol or saline, mice were placed in the apparatus and confined to the appropriate side of the chamber for 5 min. On the 10th day, mice were not treated with any drug and were tested for preference by placing them in the apparatus with access to both sides of the chamber for 30 min. The preference score was calculated as time spent on the ethanol paired side post-conditioning minus pre-conditioning.

**Supplementary Table 1.** Primer sequences for qPCR. Primers are shown in the 5' to 3' direction.

| Gene | Forward primer | Reverse primer |
|---|---|---|
| *Gusb* | CGGGACTTTATTGGCTGGGT | CCATTCACCCACACAACTGC |
| *Rpl13a* | TACCAGAAAGTTTGCTTACCTGGG | TGCCTGTTTCCGTAACCTCAAG |
| *Hprt* | GTTGGGCTTACCTCACTGCT | TCATCGCTAATCACGACGCT |
| *Gfap* | ATCGAGATCGCCACCTACAG | CTCACATCACCACGTCCTTG |
| *Slc1a2* | CGGGAAGAAGAACGACGAGG | TGACCGCCTTGGTGGTATTG |
| *Slc1a3* | CATGTGCTTCGGTTTCGTGA | TCCCTGCGATCAAGAAGAGG |
| *Slc7a11* | TGGACGCTACATCCTGGAAC | AAATCTGGATCCGGGCACTC |
| *Slc17a8* | AGCCACGACTTCCTTCTGTG | TACTGCACCAATACCCCTGC |

**Supplementary Table 2.** Statistical test results.

| Figure | Measure | Statistical Test | Result | Multiple comparisons test | Results |
|---|---|---|---|---|---|
| **1B** | Number of Aldh1l1+ cells (BaseScope) | Unpaired t-test with Welch's correction | t = 0.4690, df = 7.205<br>p = 0.6529 | N/A | None |
| **1C** | % Aldh1l1+ cells that are Stat3+ (BaseScope) | Unpaired t-test with Welch's correction | **t = 4.422, df = 9.885**<br>**p = 0.0013 **** | N/A | None |
| **1D** | Number of Stat3+ Aldh1l1– cells (BaseScope) | Unpaired t-test with Welch's correction | t = 1.445, df = 9.999<br>p = 0.1789 | N/A | None |
| **2A** | *Gfap* relative expression (qPCR) | Two-way ANOVA (Genotype x Sex) | **Genotype: F(1,43) = 68.28, p < 0.0001 ******<br>Sex: F(1,43) = 2.381, p = 0.1301<br>Interaction: F(1,43) = 0.0056, p = 0.9408 | Not performed | None |

| Figure | Measure | Statistical Test | Result | Multiple comparisons test | Results |
|---|---|---|---|---|---|
| **2B** | *Slc1a2* relative expression (qPCR) | Two-way ANOVA (Genotype x Sex) | **Genotype: F(1,45) = 7.800, p = 0.0076 ****<br>Sex: F(1,45) = 0.0535, p = 0.8181<br>Interaction: F(1,45) = 0.3219, p = 0.5733 | Not performed | None |
| **2C** | *Slc1a3* relative expression (qPCR) | Two-way ANOVA (Genotype x Sex) | Genotype: F(1,44) = 3.370, p = 0.0732<br>Sex: F(1,44) = 0.1536, p = 0.6970<br>Interaction: F(1,44) = 0.4233, p = 0.5187 | Not performed | None |
| **2D** | *Slc7a11* relative expression (qPCR) | Two-way ANOVA (Genotype x Sex) | Genotype: F(1,44) = 0.4368, p = 0.5121<br>Sex: F(1,44) = 0.0207, p = 0.8861<br>**Interaction: F(1,44) = 4.891, p = 0.0322 *** | Šídák's multiple comparisons test | Male, Cre - vs Cre +: diff = 0.096, p = 0.4528, ns<br>Female, Cre - vs Cre +: diff = –0.178, p = 0.1099, ns |
| **2E** | *Slc17a8* relative expression (qPCR) | Two-way ANOVA (Genotype x Sex) | **Genotype: F(1,44) = 7.111, p = 0.0107 ***<br>Sex: F(1,44) = 0.0198, p = 0.8888<br>Interaction: F(1,44) = 2.209, p = 0.1443 | Not performed | None |
| **3B** | GLT-1 monomer, males (WB) | Unpaired t-test with Welch's correction | t = 0.4577, df = 6.553,<br>p = 0.6620 | N/A | N/A |
| **3C** | GLT-1 dimer, males (WB) | Unpaired t-test with Welch's correction | **t = 3.204, df = 7.180,<br>p = 0.0145 *** | N/A | N/A |
| **3E** | GLT-1 monomer, females (WB) | Unpaired t-test with Welch's correction | t = 0.8283, df = 4.244,<br>p = 0.4515 | N/A | N/A |
| **3F** | GLT-1 dimer, females (WB) | Unpaired t-test with Welch's correction | t = 0.7747, df = 6.805,<br>p = 0.4646 | N/A | N/A |
| **4B** | sEPSC frequency | Unpaired t-test | t(51) = 0.1493, p = 0.8819 | N/A | N/A |
| **4C** | sEPSC amplitude | Unpaired t-test | **t(51) = 2.836, p = 0.0065**** | N/A | N/A |

| Figure | Measure | Statistical Test | Result | Multiple comparisons test | Results |
|---|---|---|---|---|---|
| **4D** | sEPSC rise time | Unpaired t-test | t(51) = 1.150, p = 0.2554 | N/A | N/A |
| **4E** | sEPSC decay time | Unpaired t-test | t(51) = 1.364, p = 0.1785 | N/A | N/A |
| **4F** | sEPSC area | Unpaired t-test | t(51) = 0.861, p = 0.3933 | N/A | N/A |
| **5A** | Male ethanol intake, DID (12 sessions) | Mixed-effects model (REML) | **Time: F(11,186) = 18.04, p < 0.0001 ****** **Genotype: F(1,17) = 10.95, p = 0.0041 **** Time x Genotype: F(11,186) = 0.5259, p = 0.8842 | Not performed | None |
| **5B** | Male sucrose intake (4 sessions) | Mixed-effects model (REML) | Time: F(2.502,54.22) = 2.820, p = 0.0568 Genotype: F(1,22) = 0.7428, p = 0.3980 **Time x Genotype: F(2.502,54.22) = 3.154, p = 0.0401 *** | Not performed | None |
| **5C** | Male blood ethanol clearance | Two-way RM ANOVA | **Time: F(2.274, 20.46) = 146.5, p<0.0001****** Genotype: F(1, 9) = 2.514, p=0.1473 Time x Genotype: F(2.274, 20.46) = 0.5962, P=0.5807 | Not performed | None |
| **5D** | Female ethanol intake, DID (12 sessions) | Two-way RM ANOVA | **Time: F(11,187) = 15.54, p < 0.0001 ****** Genotype: F(1,17) = 0.0523, p = 0.8218 Time x Genotype: F(11,187) = 0.6699, p = 0.7658 | Not performed | None |
| **5E** | Female sucrose intake (4 sessions) | Two-way RM ANOVA | Time: F(3,51) = 1.792, p = 0.1605 Genotype: F(1,17) = 0.3740, p = 0.5489 Time x Genotype: F(3,51) = 1.056, p = 0.3761 | Not performed | None |
| **5F** | Female blood ethanol clearance | Two-way RM ANOVA | **Time: F(1.994, 19.94) = 62.56, P<0.0001****** Genotype: F(1, 10) = 0.04513, p = 0.8360 Time x Genotype: F(1.994, 19.94) = 0.6191, p = 0.5480 | Not performed | None |

**Supplementary Table 3: Putative STAT3 Binding Sites in *Slc1a2***

FIMO scan ($p < 0.0001$) using STAT3 motif MA0144.1 (JASPAR). Genome assembly: mm10. Gene: Slc1a2, chr2, + strand, TSS = 102,658,659 (GENCODE ENSMUST00000080210.9). Distance formula: site center − TSS. Negative = upstream (promoter). Positive = downstream (gene body). Sites ordered by distance from TSS.

| Parameter | Site 2 | Site 4 | Site 3 | Sites 1+5 * |
|---|---|---|---|---|
| **Genomic coordinates (mm10)** | chr2:102,658,531–541 | chr2:102,658,961–971 | chr2:102,660,087–097 | chr2:102,664,124–134 * |
| **Strand** | – | + | – | +* |
| **Distance to TSS** | −123 bp | +307 bp | +1,433 bp | +5,470 bp |
| **Location** | Proximal promoter | Exon 1 (5′UTR) | First intron | First intron |
| **Matched sequence** | TTCTTGGAAG | TTCAGGAAGG | TTCCCAGAAG | TTCCTGGAAG * |
| **FIMO p-value** | $2.41 \times 10^{-5}$ | $3.81 \times 10^{-5}$ | $2.87 \times 10^{-5}$ | $1.24 \times 10^{-5}$ |

* Sites 1 and 5 represent a single palindromic STAT3 binding element (GAS motif). Site 5 (chr2:102,664,123–133, strand –, $p = 9.08\times10^{-5}$) and Site 1 (chr2:102,664,124–134, strand +, $p = 1.24\times10^{-5}$) overlap by 9 of their 10 bp and share the same center position. FIMO detected the near-palindromic STAT3 consensus (TTCnnnGAA) on both strands at the same locus. These are reported here as a single functional site using the more significant p-value ($p = 1.24\times10^{-5}$, Site 1).

**Supplementary Table 4.** Effect of genotype on intrinsic membrane properties in medial division CeA neurons of Cre – (control) and Cre + (Stat3 aKO) mice. Recordings were performed with K-gluconate -based internal solution. Between group comparisons by two-tailed t-test, *p<0.05. The data are expressed as the mean ± SEM.

| | **Cre –** (n=38) | **Cre +** (n=44) | **Statistics** |
|---|---|---|---|
| **Vrest (membrane resting potential, mV)** | -59.93 ± 0.62 | -59.52 ± 0.64 | t=0.4476, df=80<br>p = 0.656 |
| **Rmax (membrane resistance, MΩ)** | 494.2 ± 18.46 | 489.5 ± 23.79 | t=0.1551, df=80<br>p = 0.877 |
| **Time constant (membrane time constant, ms)** | 51.74 ± 2.93 | 55.96 ± 3.03 | t=0.9993, df=77<br>p = 0.321 |
| **Capacitance (membrane capacitance, pF)** | 66.12 ± 1.97 | 70.13 ± 2.96 | t=1.128, df=74<br>p = 0.263 |
| **Membrane voltage sag slope (mV/nA)** | -116.3 ± 12.24 | -107.1 ± 9.70 | t=0.5960, df=79<br>p = 0.553 |
| **Afterdepolarization slope (mV/nA)** | -162.6 ± 24.21 | -118.4 ± 12.62 | t=1.680, df=56<br>p = 0.099 |

**Supplementary Table 5.** Effect of genotype on AP characteristics in medial division CeA neurons of Cre – (control) and Cre + (Stat3 aKO) mice. Between group comparisons by two-tailed t-test, *p<0.05. The data are expressed as the mean ± SEM.

| | **Cre –** (n=39) | **Cre +** (n=44) | **Statistics** |
|---|---|---|---|
| **Rheobase (pA)** | 19.28 ± 2.58 | 19.82 ± 2.28 | t=0.16, df=62<br>p = 0.874 |
| **Threshold (mV)** | -38.53 ± 0.58 | -38.58 ± 0.62 | t=0.06, df=79<br>p = 9.950 |
| **Amplitude (mV)** | 84.06 ± 1.28 | 84.14 ± 1.02 | t=0.05, df=80 |

| | | | |
|---|---|---|---|
| | | | p = 0.958 |
| **Half-width (mV)** | 2.020 ± 0.05 | 2.132 ± 0.07 | t=1.20, df=77<br>p = 0.233 |
| **Kink** | 254.9 ± 15.60 | 271.8 ± 11.91 | t=0.87, df=81<br>p = 0.385 |
| **Kink slope** | 0.978 ± 0.12 | 0.909 ± 0.12 | t=0.41, df=79<br>p = 0.686 |
| **Upshoot slope (Vm/ms)** | 115.7 ± 5.51 | 118.7 ± 4.28 | t=0.45, df=81<br>p = 0.656 |
| **Spike positive phase** | 0.619 ± 0.01 | 0.611 ± 0.01 | t=1.00, df=79<br>p = 0.322 |
| **Decay slope (Vm/ms)** | -40.64 ± 1.73 | -42.10 ± 2.00 | t=0.55, df=81<br>p = 0.586 |
| **Spike negative phase** | 0.742 ± 0.01 | 0.745 ± 0.01 | t=0.25, df=81<br>p = 0.803 |

### Supplementary Figures

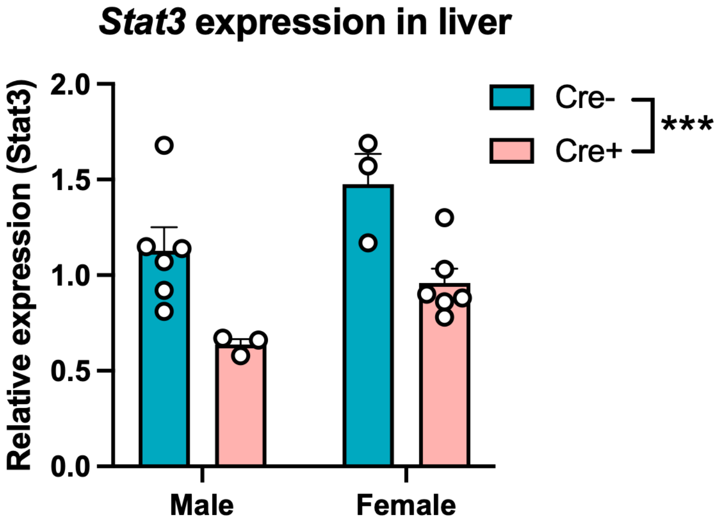


**Supplementary Figure 1. Reduced *Stat3* expression in the liver of Stat3 aKO mice.** RNA was isolated from the livers of Stat3 aKO (Cre +, n=3 male and 6 female) and control Stat3$^{flox}$ (Cre -, n=6 male and 3 female) mice and cDNA was synthesized and subjected to qPCR with *Stat3* primers. ***p<0.001, main effect of genotype ($F_{(1, 14)}$ = 18.33, p = 0.0008) and *p<0.05, main effect of sex ($F_{(1, 14)}$ = 8.066, p = 0.0131) by two-way ANOVA.

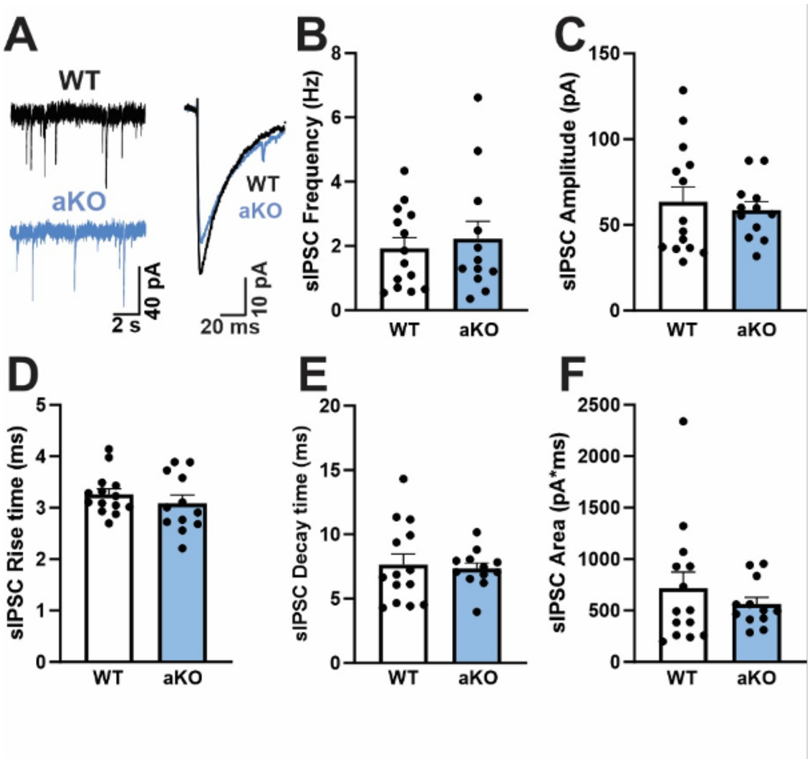


**Supplementary Figure 2. Astrocyte *Stat3* deletion does not affect baseline spontaneous GABA transmission in the central nucleus of the amygdala. (A**) Representative traces of sIPSCs from a CeA neuron (left panel) and average amplitude of sIPSC (right panel) of wild-type and Stat3 aKO mice. (B) Baseline frequency. (C) Amplitude, (D) rise time, (E) decay time and (F) area of sIPSCs in Stat3 aKO (Cre +; n = 12 cells) and Stat3 flox (Cre -; n = 14 cells) mice. The data are presented as mean ± SEM.

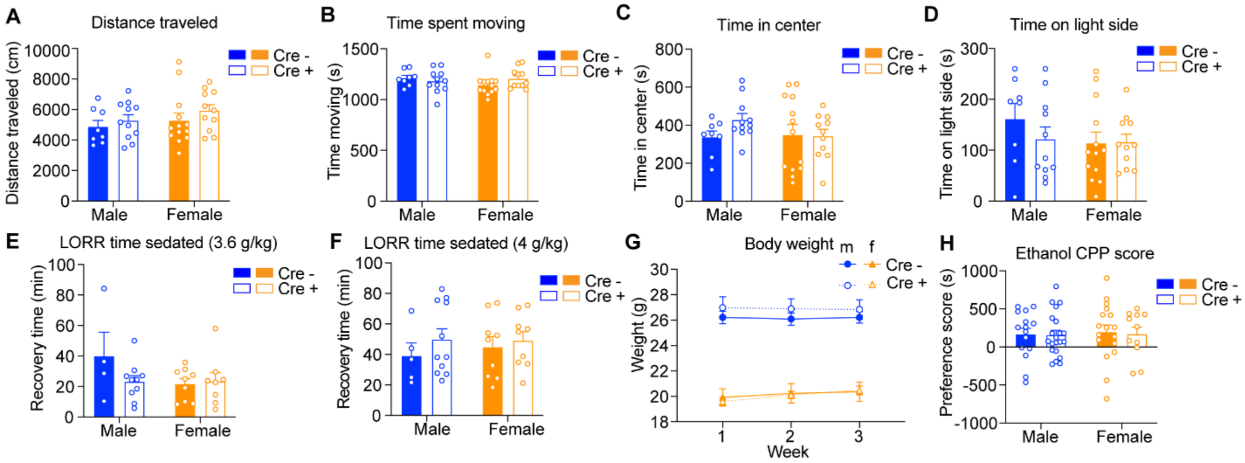


**Supplementary Figure 3. Astrocyte *Stat3* deletion does not affect baseline locomotor activity, anxiety-like behavior or ethanol-associated behaviors.** (**A**) Distance traveled in an open field in Cre - (8 males and 13 females) and Cre + (11 males and 11 females) mice. (**B**) Time spent moving in the open field in Cre - and Cre + mice. (**C**) Time spent in the center of the open field in Cre - and Cre + mice. (**D**) Time spent on the light side of a light-dark box in Cre – (8 males and 13 females) and Cre + (11 males and 11 females) mice. The same mice were used for the open field and light-dark box test. (**E**) Sedation time in the ethanol loss-of-righting reflex (LORR) test after 3.6 g/kg i.p. ethanol injection in Cre – (4 males and 9 females) and Cre + (10 males and 8 females) mice. (**F**) Sedation time in the ethanol LORR test after 4 g/kg i.p. ethanol injection in Cre – (5 males and 9 females) and Cre + (11 males and 9 females). The same mice were used for 3.6 g/kg and 4 g/kg LORR and tested one week apart. The number of mice varies between doses due to mice being excluded for not sedating after ethanol injection (1 Cre – and 2 Cre + males and 1 Cre + female at 3.6 g/kg; 1 Cre + male at 4 g/kg). (**G**) Weekly body weights of male and female Cre – and Cre + mice during the ethanol drinking in the dark test. (**H**) Ethanol conditioned place preference (CPP) score in Cre – (15 males and 17 females) and Cre + (21 males and 11 females) mice. Data are graphed as the mean ± SEM.
